# U-Net Ensembles for Segmentation of High-Density Cell Populations and Organelles

**DOI:** 10.1101/2024.11.19.623228

**Authors:** Samuel J. Fulton, Cameron Baenen, Irina D. Pokrovskaya, Maria A. Aronova, Brian Storrie, Richard D. Leapman

## Abstract

The complex and highly intertwined morphology of activated platelets within blood clot thrombi poses significant challenges for segmentation. Here we develop a robust machine vision pipeline for cell and organelle segmentation within volume electron microscope (vEM) datasets and validate that against both model CREMI neuron segmentation challenge data and focused isotropic FIB-SEM imaged platelets and their organelles within a series of ferric chloride indued murine occlusive clots Neural network predictions were collected along multiple planes, capturing 3D correlations using only 2D neural networks. Using a workstation class computer, we segmented and analyzed hundreds of platelets and report quantitative morphological measurements of platelets and their thousands of included organelles. CREMI neuron segmentation challenge analysis produced state-of-the-art outcomes as did platelet data in comparison to manual segmentation. Our work paves the way for large-scale, single cell, 3D studies across multiple examples and lends initial single cell level insights into occlusive thrombus structure and platelet heterogeneity therein. We include a lightweight Jupyter notebook for initializing and running the neural network (Data and Materials Availability). We also provide Amira Avizo protocols for converting neural network outputs to cell and organelle instances and subsequent morphological measurements.

## Introduction

Recent advances in high-throughput volume electron microscopy (vEM) have made it possible to acquire giga voxel-scale datasets within just a matter of hours. In stark contrast, extracting biological insights from these datasets demands an extensive investment of human labor, often amounting to thousands of human hours [Guay et al. 2021; Alwath et al., 2023; Berning et al.,2015]. The emergence of deep learning methodologies has demonstrated remarkable efficacy in automating the traditionally labor-intensive process of manual segmentation [Helmstaedter et al, 2013; Zhou et al., 2018; Schlemper et al., 2019; Ibtehaz and Rahman, 2020; Plaza and Funke, 2018]. Nonetheless, the 3D segmentation of densely packed cells using a single stack of 2D image planes poses a challenge due to ambiguous membrane boundaries having irregular and complex shapes. This is exemplified by failure to properly segment a platelet when part of its membrane surface happens to be oriented parallel to an image plane of the acquired volume.

To address this, we developed a multi-axis analysis pipeline that aggregates 2D convolutional neural network predictions across three orthogonal planes to generate accurate 3D segmentations. Our approach integrates two complementary U-Net–based networks: one trained to detect platelet membranes and another trained to estimate distance from membranes. By linearly combining their predictions, we produce a topographical landscape that can be efficiently partitioned into distinct cell instances using a watershed clustering algorithm. In doing so, we improve segmentation performance and provide a state-of-the-art solution that serves as a practical, workstation-level alternative to computationally intensive 3D neural networks.

We validated segmentation performance against the CREMI Serial Block Face Scanning Electron Microscopy (SBF-SEM) neuron dataset [Funke et al., 2016], achieving strong accuracy despite the dataset’s anisotropic resolution. We then benchmarked our pipeline against ilastik [Berg et al., 2019], representing a classical semi-automated baseline commonly used in cell biology, and U-Net [Isensee et al., 2021], representing a modern deep learning baseline. Finally, we applied the pipeline to isotropic FIB-SEM Focused Ion Beam Scanning Electron Microscopy datasets of platelet-rich thrombi formed in mouse carotid arteries under varying inducer conditions, as well to a resting platelet dataset as control. This enabled quantitative analysis of thousands of platelets and organelles, including mitochondria, α-granules, and dense granules.

From the observed morphologies, we were able to infer underlying structural dynamics and identify distinct patterns of platelet remodeling. We found that dense granules showed extensive depopulation relative to other organelles, whereas α-granules (both condensed and their substantially released decondensed form [Pokrovskaya et al., 2016; Pokrovskaya et al., 2018]) and mitochondria exhibited binary depletion behavior in which a subpopulation of platelets exhibited a complete loss of granules and mitochondria across all inducer strengths. Moreover, we observed evidence for highly local clustering and short-range degranulation events, providing a preliminary challenge to prevailing models of thrombus formation as a process emanating outward in uniform gradients from a damage site.

Together, this work establishes both a methodological advance, being a practical, readily transferable framework for large-scale 3D cellular segmentation, and a biological advance that reveals, at an unprecedent scale, novel examples of platelet activation state at the 3D, single cell level in occlusive clots

## Materials and Methods

### Experimental Model and Study Participant Details

Wild-type C57BL/6 mice (male and female, 8–12 weeks old) were obtained from Jackson Laboratories (Bar Harbor, ME) and used in equal numbers. All animal care complied with and received approval (#A3063-01) from the Institutional Animal Care and Use Committee of the University of Kentucky. Two independent mouse experiments were performed at each ferric chloride concentration.

## Method Details

### Thrombus Induction and Sample Preparation

Carotid artery thrombi were generated in wild-type mice by topical application of ferric chloride (FeCl₃) to the external vessel wall, a standard method for inducing endothelial injury and occlusive thrombosis [Joshi et al., 2023]. This procedure was performed by the Whiteheart laboratory at the University of Kentucky. Following thrombus formation, arterial segments containing the occlusion were fixed in situ, excised, post fixed ex vivo, and sent to the Storrie laboratory at the University of Arkansas for Medical Sciences (UAMS) for additional fixation, washing, heavy metal staining, resin embedding, and trimming. Samples were then shipped to the Leapman laboratory at the NIH for SBF- and FIB-SEM imaging. All samples were SBF-SEM imaged in duplicate (2 independent mouse experiments) before FIB-SEM ROI analysis.

Unactivated (resting) platelet samples were prepared from whole blood drawn from healthy mice and immediately mixed with fix. Platelets were isolated by differential centrifugation and suspended in a resting buffer to preserve their quiescent state prior to fixation and processing (Pokrovskaya et al,216).

#### Electron Microscopy

Resin-embedded samples were mounted and coated with thin layers of carbon and gold to enhance conductivity and image quality. Regions of interest (ROIs) were imaged using a Zeiss Crossbeam 550 FIB-SEM (Carl Zeiss Microscopy, LLC) under high vacuum, with an accelerating voltage of 1.5 kV, beam current of 2 nA, and a 55 µm condenser aperture. For milling, the ion beam operated at 30 kV, with currents of 700 pA for shallow sections, increased to ≥1.5 nA for sections deeper than 20 µm. Beam conditions were optimized per dataset to minimize curtaining and preserve ultrastructural detail. Datasets were collected at near-isotropic voxel sizes of 5 nm × 5 nm × 7 nm with dwell times of ∼10 µs. System calibration and alignment were performed before each acquisition, which was managed using Atlas Engine and Atlas 5 software (Fibics Inc., Ottawa, Canada).

Whole-clot structure was first imaged using SBF-SEM at a voxel size of 160 nm × 160 nm x 1000 nm to provide a lower resolution, large-volume overview of thrombus architecture. At this resolution, fine ultrastructural features are not resolved; however, mesoscale organization, including platelet-rich regions, red blood cell-rich areas, and variations in clot density and morphology, can be readily distinguished. These features were used to guide selection of ROIs representing a range of spatial locations and microenvironments across the clot (e.g., core versus periphery and regions of differing cellular composition).

The sample was then transferred to the FIB-SEM where high-resolution datasets (<10 nm voxel size; maximum ∼40 μm × 40 μm× 40 µm) were then acquired in selected ROIs. Due to the destructive nature of SBF-SEM, FIB-SEM imaging was not performed on the same physical volume previously imaged by SBF-SEM, and no direct volumetric registration between modalities was attempted. Instead, SBF-SEM and FIB-SEM were used in a complementary, multiscale manner rather than as a strictly correlative workflow imaging the exact same physical volume. SBF-SEM served as a screening step to identify regions of interest, while FIB-SEM enabled detailed characterization of subcellular ultrastructure within those regions.

This multiscale imaging workflow is inherently time intensive. Acquisition of SBF-SEM data to approximately the midpoint of the clot required ∼1 week of imaging. High-resolution FIB-SEM acquisition of individual ROIs required approximately 1–2 days per volume, with ∼8 ROIs typically sampled per thrombus. Following FIB-SEM imaging, SBF-SEM acquisition was resumed to complete the volume, requiring an additional ∼1 week. These acquisition times underscore the substantial experimental demands associated with large-volume, high-resolution volumetric electron microscopy of thrombi. 3D images were rendered for 3 representative ROIs at each ferric chloride concentration. This integrated approach captures the hierarchical organization of the clot, linking nanoscale ultrastructure to global architecture.

#### Neural network architecture

We employed a single-channel convolutional neural network due to the lack of overlapping hand-labeled cell membrane and organelle data. The network architecture is based on the U-Net framework, originally introduced by Ronneberger et al. in 2015; Siddique et al., 2021), which has since been widely used in biomedical imaging [Zhou et al., 2018; Schlemper et al.m 2019; Ibtehaz and Rahman, 2020].

The encoder consists of an initial convolutional layer followed by four residual blocks derived from ResNet18. The bottleneck contains two convolutional layers with ReLU activations. The decoder consists of three convolutional and up sampling blocks, and three residual connections are incorporated to stabilize training. A final 1 × 1 convolution layer with sigmoid activation produces the segmentation probability maps and distance maps.

#### U-Net training

The segmentation pipeline utilizes two complementary networks trained on membrane annotations. Images were manually labeled using Amira Avizo software, and corresponding distance maps were generated using Amira’s “distance map” module. The labeled dataset was split into 90% training and 10% validation data.

Data augmentation was applied to increase dataset diversity, including elastic deformations, rotations, brightness and contrast adjustments, and Gaussian noise. Each image was randomly cropped into 100 patches of 1024 × 1024 pixels, each receiving a random combination of augmentations. Networks were trained for 20 epochs with a batch size of 32, using early stopping with a patience of five epochs. Optimization was performed using the Adam optimizer with a learning rate of 1 × 10⁻⁵ and L2 regularization. Binary cross-entropy loss was used for membrane probability map generation.

The proposed multi-axis segmentation framework is computationally efficient and can be trained on modest hardware. Training was conducted on a single NVIDIA A100 GPU on the NIH Biowulf computing cluster, requiring approximately 30 minutes per network. We also trained using a local consumer-grade GeForce GTX 1080 GPU which required a longer training time of two hours. Because the approach operates on 2D slices rather than full 3D volumes, memory requirements are substantially reduced compared to 3D U-Net architectures, which typically require significantly greater GPU memory and longer training times. This enables rapid training and iteration, as well as deployment on widely accessible hardware. The multi-axis strategy provides a practical compromise, capturing 3D contextual information through aggregation of orthogonal 2D predictions while avoiding the computational overhead associated with fully 3D models.

### Post-processing and feature extraction

Following neural network segmentation, masks were imported into Fiji [Schindelin et al., 2012] and Amira 3D 2023.2 (Thermo Fisher Scientific) for post-processing. Instance-level segmentations were refined using watershed-based separation, inspected to correct obvious artifacts, and rendered in 3D for visualization. The Border Kill module in Amira was used to retain only segmented cells fully contained within the ROI. Using Amira’s Label Analysis module, per-instance measurements (e.g., volume, surface area, centroid position) were extracted for platelets and organelles. These data were then exported for downstream statistical analysis in Python.

### Quantification and Statistical Analysis

Statistical analyses were performed using Python (v3.12.7) with the scipy.stats and statsmodels libraries. To compare fractional organelle volumes across experimental conditions, we first excluded platelets lacking the organelle of interest (i.e., cells with a fractional volume of zero), focusing our analysis on cells that retained granules or mitochondria. Group comparisons were then assessed using the Kruskal-Wallis H test, a non-parametric method appropriate for comparing more than two independent groups without assuming normal distributions. A significance threshold of *p* < 0.05 was used. In cases where the Kruskal-Wallis test revealed significant differences, post hoc pairwise comparisons were planned with multiple testing correction, though not needed for the final organelle volume analysis due to a non-significant result.

Sample sizes were defined as the number of individual platelets analyzed per condition. The immediately fixed control dataset contained 77 platelets. For the 5.5% occlusion condition, ROIs 1–3 contained 39, 134, and 96 platelets, respectively. The 6% condition included 34, 32, and 44 platelets across ROIs 1–3 (due to taking smaller volumes of FIB-SEM data), and the 10% condition included 115, 118, and 172 platelets across ROIs 1–3.

## Results

### Developments and summary of segmentation pipeline

Deep learning segmentation methodologies have advanced rapidly, encompassing supervised, semi-supervised, and unsupervised paradigms [Aswaath et al., 2023]. Fully supervised 3D segmentation techniques have achieved state-of-the-art performance, enabling cellular analysis at scales previously deemed infeasible [Heinrich et al., 2021; Shapson-Coe et al., 2024; Jia et al., 2020]. Weakly supervised and unsupervised techniques have likewise proven effective in settings where annotated training data are limited [Wolney et al., 2021; Ma et al., 2020; Bermudez-Chacon er al., 2020]. More recently, foundation models such as Segment Anything and emerging 3D extensions (SAM 3D) have demonstrated strong performance in prompt-based segmentation tasks [SAM3D Team, 2025].

The conventional approach to instance-level segmentation typically employs a U-Net architecture to generate probability maps, which indicate the likelihood of a pixel belonging to a specific object. These maps are subsequently processed using classical algorithms, such as watershed clustering, to achieve instance-level segmentation [Ma et al., 2020, Cireşan, et al., 2012; Turaga et al., 2010; Shen, 2017; Matharu et al, 2023; Franco-Barranco et al., 2022].

However, pixel classification methods often suffer from the erroneous splitting of large cells. To mitigate this limitation, alternative approaches have been developed wherein neural networks are trained to predict distance metrics from the background. Euclidean distance-based methods have shown success in reducing cell fragmentation [Ma et al., 2020, Wang et al., 20219; Schmidt et al. 2018; Faza et al., 2019; Zhou et al., 2019; Naylor et al., 2019; Spierrs et al., 2021] but can misidentify pseudopods and other branching structures at cell surfaces. More critically, unlike many biological datasets where cells are spatially separated, platelets within thrombi are tightly interwoven, with membranes often directly apposed. This crowded ultrastructural context makes conventional deep learning segmentation approaches prone to failure, highlighting the need for more robust strategies.

To address these limitations, we integrate membrane identification with distance mapping, thereby combining the strengths of both approaches **(Fig. 1)**. To further enhance segmentation accuracy, we take advantage of the isotropic 3D nature of FIB-SEM data by rotating the volume and aggregating 2D network predictions across multiple axes. In many vEM datasets, limited z-resolution necessitates cross-slice attention models that mimic radiologists’ reasoning [Heinrich et al., 2021; Al-Kofahi et al., 2018; Hung et al., 2024; Huang et al., 2021]. In contrast, the isotropic nature of FIB-SEM datasets enables a more efficient solution: by projecting 3D correlations from multi-axis 2D predictions, we achieve high segmentation accuracy without the computational cost of fully 3D networks [Spiers et al., 2021; Stringer et al., 2021], and therefore a workstation class computer can be sufficient.

**Fig. 1.**
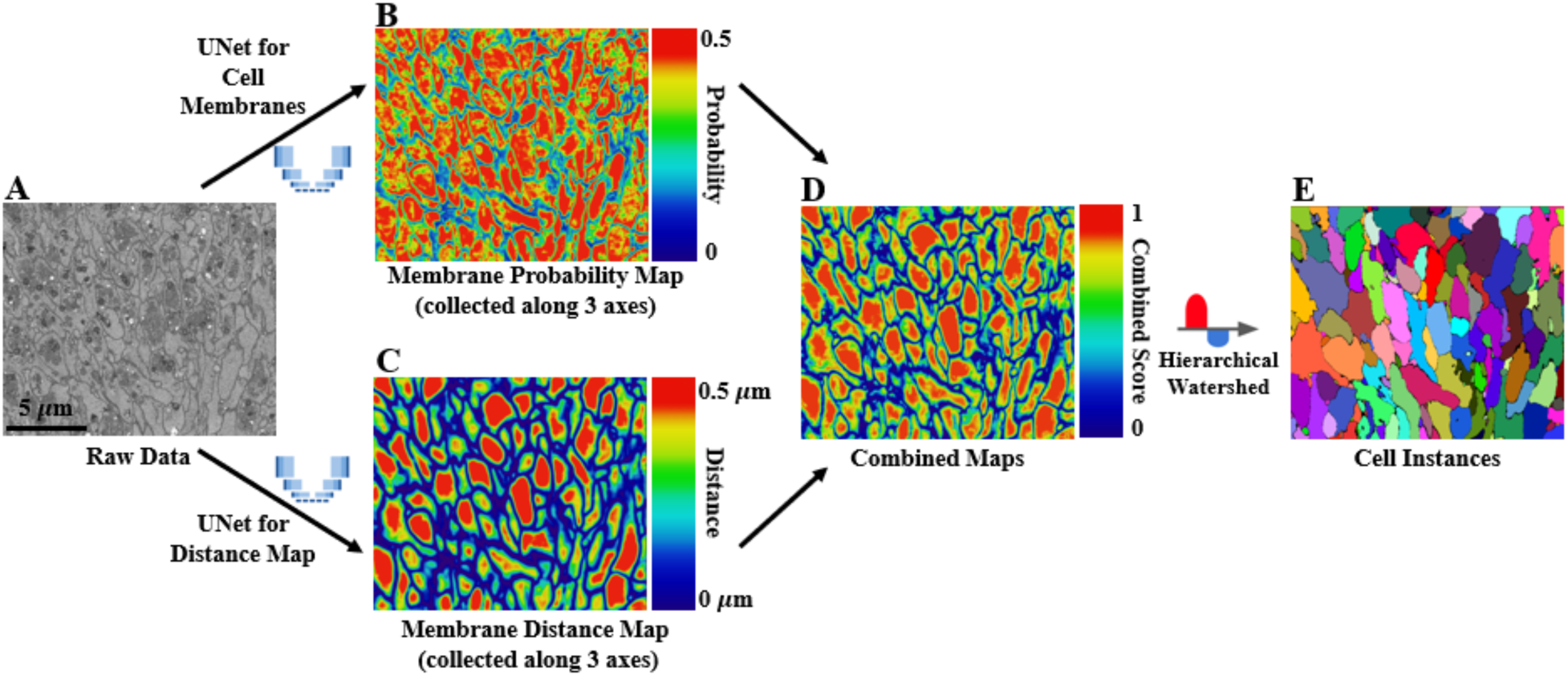
Segmentation pipeline. **(A)** Raw electron microscope data, 5 nm x5 nm x7nm voxel size. **(B)** Membrane probability map, averaged from multi-axis analysis. **(C)** Predicted distance to the nearest membrane, averaged from multi-axis analysis. **(D)** Normalized combination of (B) and (C) (scaled to one and averaged). **(E)** Final cell instances generated using H-extrema watershed. Note that the same workflow can be applied to segment other classes of interest, such as mitochondria or α-granules.

The development of this field has been significantly supported by publicly available FIB-SEM datasets, such as Lucchi++ (mouse hippocampus), FIB-25 (drosophila optic lobe), Open Organelle (Hela cells), Cardiac Mitochondria (mouse heart), and UroCell (mouse urothelial cells) [Aswath et al., 2023]. Building on this foundation, we analyze a novel FIB-SEM dataset of platelet-rich thrombi in mouse carotid arteries, providing new insights into clot structure at nanometer resolution. Our work advances platelet segmentation from the semantic segmentation of Guay et al. [2021] to full instance-level segmentation. By focusing on platelets within intact thrombi, we extend beyond the analysis of isolated platelets performed by Matharu et al. [2023]. To our knowledge, this represents the first large-scale 3D segmentation of individual platelets within thrombi, marking a significant step forward in both methodology and application.

In parallel, though not discussed explicitly in this paper, our labs are developing 2D activation state maps that register segmented regions of interest (ROIs) within the broader thrombus cross-section [Faruque et al., 2025] These complementary efforts will enable integration of single-platelet remodeling with global thrombus organization.

### Model benchmarking and evaluation

First, we benchmarked our whole-cell segmentation pipeline using the CREMI neuron segmentation challenge dataset, which consists of 125 hand-labeled slices of 1250 × 1250 pixels each. Unlike our near-isotropic FIB-SEM dataset, the CREMI data are highly anisotropic, with voxel dimensions of 4 nm × 4 nm × 40 nm. To utilize multi-plane analysis, we trained two neural networks. The first network was trained on 12 slices in the xy-plane. For the second network, we resampled these slices to generate 120 slices with pixel dimensions of 4 nm × 40 nm, matching the xz and yz planes. A slice of our segmentation results is shown alongside the raw data in **Figure 2**.

**Fig. 2.**
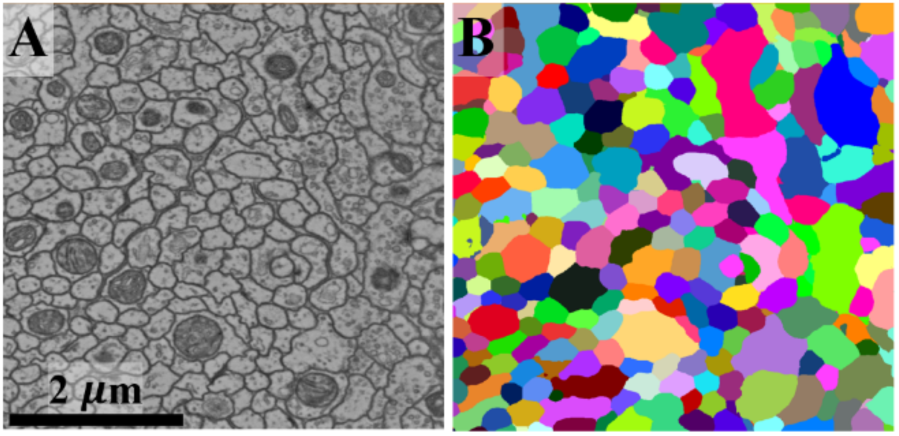
Benchmark segmentation results. **(A)** Raw SBF-SEM neuron data from the CREMI segmentation challenge. **(B)** Whole-cell segmentation results, where each color represents a unique cell (colors may be reused).

Segmentation was benchmarked by classifying pixels near cell boundaries as membrane and comparing them with the ground truth. We computed precision, recall, accuracy, and F1 scores **(Table I)**.

**Table I:**
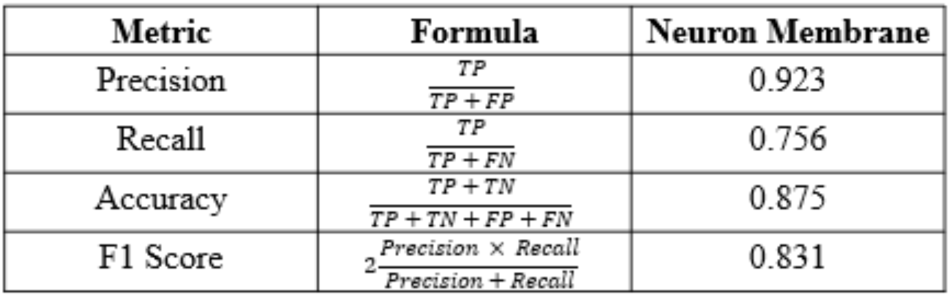
The metric formulas for segmentation are expressed in terms of True Positive (TP), True Negative (TN), False Positive (FP), and False Negative (FN).

Our pipeline demonstrates strong performance despite the dataset’s anisotropic nature, achieving a pixel-level precision of 0.923 in distinguishing densely packed neuronal structures. While the U-Net leverages high xy-resolution, watershed segmentation is limited by the low z-resolution in these anisotropic datasets [Meyer et al., 1990; Kornilov et al.m 2022]. Although TASCAN [Wang et al., 2022] has been proposed to improve watershed performance under anisotropy, we do not implement it here. Despite these challenges, membrane classification remains reliable, with accuracy and precision exceeding 0.85.

To illustrate the contribution of the multi-axis fusion strategy, we performed an ablation study comparing the full multi-axis pipeline to a single-axis implementation **(Fig. 3)**. In the single-axis setting, segmentation is performed independently on 2D slices, limiting contextual information along the orthogonal dimension and leading to discontinuities and slice artifacts in reconstructed 3D volumes. In contrast, multi-axis fusion integrates predictions across orthogonal planes, improving structural continuity and reducing slice-dependent artifacts. This improvement in 3D continuity is particularly important for downstream analysis, where discontinuities can propagate into errors in instance segmentation and morphological measurements.

**Figure 3.**
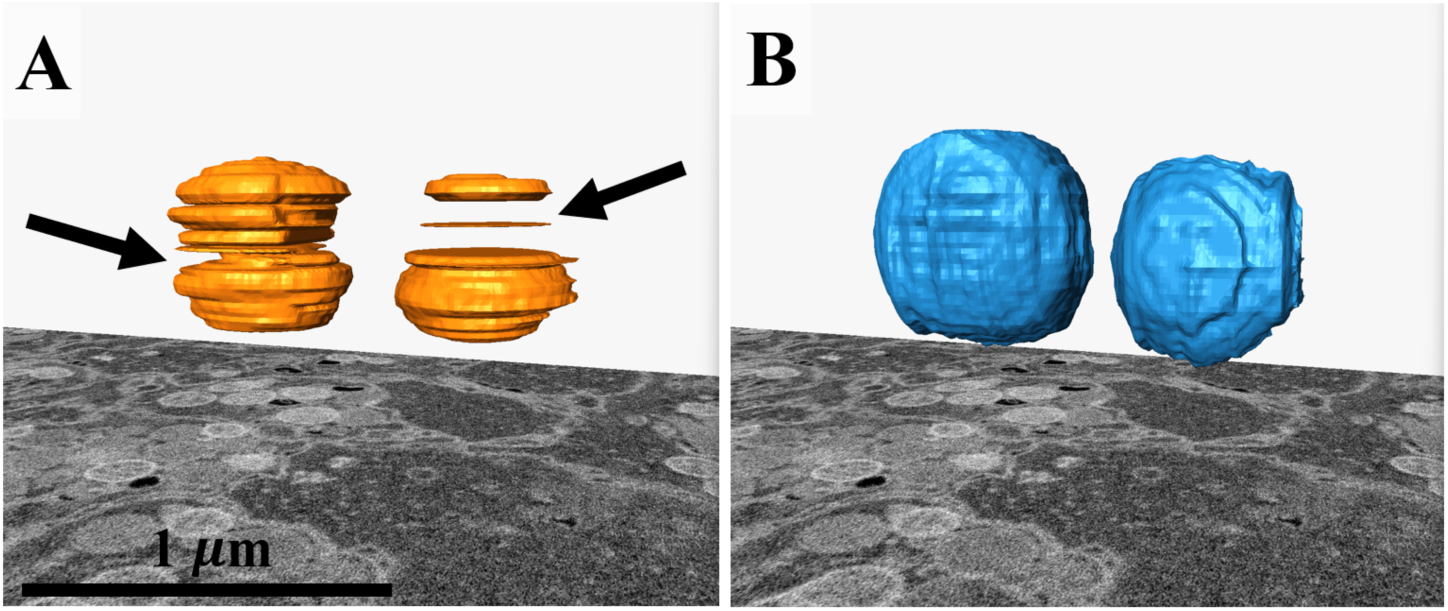
Multi-axis fusion improves segmentation by incorporating 3D contextual information. **(A)** Surface rendering of two representative mitochondria segmented using a single-axis (XY) pipeline. Because segmentation is performed independently on each slice, the resulting 3D reconstruction exhibits discontinuities, jagged surfaces, and visible slice boundaries (black arrows), reflecting limited contextual information along the z-axis. **(B)** The same mitochondria segmented using multi-axis fusion (XY, XZ, and YZ), which integrates information across orthogonal planes. This approach reduces slice artifacts and produces smoother, more continuous 3D structures. Constrained smoothing was applied to both surfaces.

Finally, to further evaluate the effectiveness of the multi-axis approach, we compared our pipeline to Ilastik and nnU-Net as representative classical semi-automated and modern deep learning baselines, respectively **(Fig. 4)**. The dataset consisted of a 1200 nm × 1200 nm × 57 μm volume of hand-labeled platelet membranes, with voxel dimensions of 5 nm × 5 nm × 7 nm. The volume was divided into four quadrants (600 nm × 600 nm × 57 μm), with three used for training and one held out for evaluation. All methods were trained and evaluated on the same data split to ensure a fair comparison. A representative XY slice from the prediction volume for each method is shown in **Figure 4**. We considered comparisons to foundation models such as SAM 3D or Cellpose-SAM; however, these approaches are primarily designed for prompt-based or interactive segmentation rather than dense, pixel-level semantic segmentation of membrane structures, making direct quantitative comparison in this context non-trivial.

**Figure 4.**
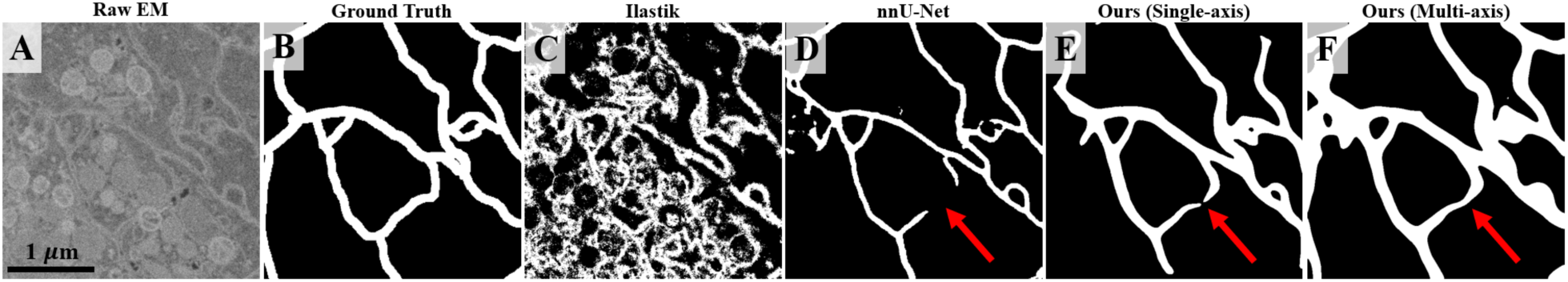
Comparison of membrane segmentation methods. **(A)** Raw EM image. **(B)** Ground truth membrane annotations. **(C)** Ilastik segmentation, which exhibits substantial noise and poor delineation of membrane boundaries. **(D)** nnU-Net segmentation, which produces high-precision predictions but fails to capture complete membrane structures, resulting in discontinuities (arrow). **(E)** Our single-axis segmentation, which improves overall accuracy but still exhibits occasional gaps in membrane continuity (arrow). **(F)** Our multi-axis segmentation, which integrates information across orthogonal planes to produce more continuous membrane predictions, resolving discontinuities observed in other methods (arrow). While this approach introduces minor over-segmentation in some regions, it substantially improves membrane completeness, which is critical for downstream watershed-based segmentation.

Membrane segmentation is inherently challenging due to the thin, sheet-like structure of membranes, where even a one-pixel deviation can substantially impact overlap-based metrics such as the Dice score. In this context, Ilastik produces noisy predictions with poor membrane delineation, while nnU-Net achieves high local accuracy but frequently fails to recover complete membrane structures, resulting in fragmented predictions. Our single-axis model reduces these errors but still exhibits occasional discontinuities. In contrast, multi-axis fusion integrates complementary information across orthogonal planes, substantially improving membrane continuity and structural completeness.

These differences are reflected quantitatively **(Table II)**. While our approach achieves competitive Dice scores, the primary improvement from the multi-axis strategy is an increase in recall (0.6974 to 0.9105), reflecting more complete membrane recovery, with a modest reduction in precision. This effect is further captured by the F2 score, which increases from 0.7035 to 0.8469, emphasizing the benefit of improved recall for this task. This tradeoff is imperative for downstream analysis: improved membrane continuity reduces the likelihood of artificial gaps that can lead to “leakage” during watershed-based instance segmentation, where adjacent regions may otherwise merge incorrectly.

**Table II:**
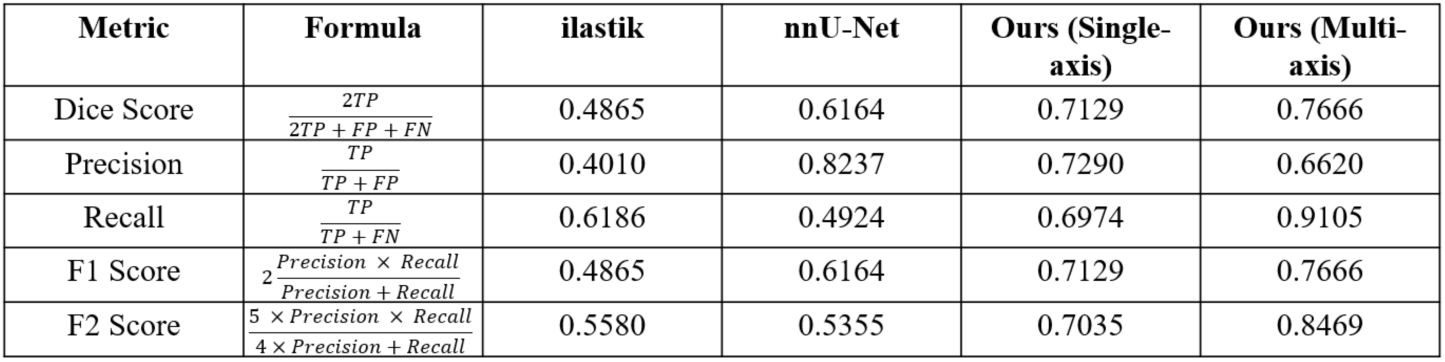
Quantitative comparison of membrane segmentation performance across methods.

### Segmentation classes in platelet samples

We performed instance-level segmentation of platelets, α-granules (condensed and decondensed), and mitochondria [Pokrovskaya et al., 2020]. Representative examples are shown in **Figure 5**. For benchmarking, we converted instance-level segmentations into semantic labels for each class. Quantitative evaluation of segmentation accuracy for the FIB-SEM platelet dataset is provided in **Table III**, reporting precision, recall, and overall accuracy. Performance was robust across all target structures, with metrics consistently above 0.85.

**Fig. 5.**
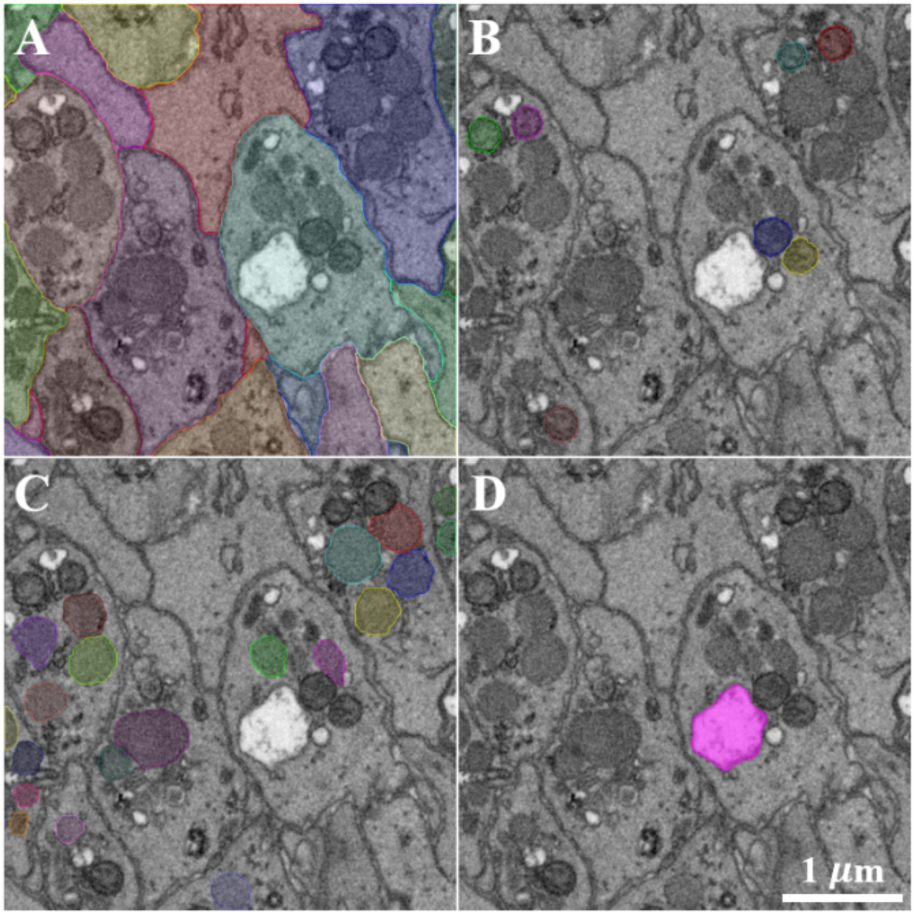
Classes of interest. Segmentation of **(A)** whole platelets, **(B)** mitochondria, **(C)** condensed α-granules, and **(D)** decondensed α-granules. Each color represents a unique instance.

**Table III:** Membrane classification demonstrates strong performances in all metrics (as defined in. **Table I).**

| Metric | Mitochondria | Alpha Granules | Platelet Membrane |
| --- | --- | --- | --- |
| Precision | 0.96 | 0.91 | 0.99 |
| Recall | 0.60 | 0.79 | 0.96 |
| Accuracy | 0.74 | 0.85 | 0.98 |
| F1 Score | 0.99 | 0.97 | 0.99 |

Segmentation of the cell plasma membrane, critical for accurate morphological measurements, achieved the highest performance, reflecting the strength of our multi-axis, membrane-focused network. Slightly reduced metrics for α-granules and mitochondria were primarily due to edge-related under-segmentation, a common challenge in high-density, crowded organelle environments. We manually segmented dense-core granules because their low abundance makes neural network prediction challenging [Müller et al., 2024]. This reliance on manual annotation for rare or difficult-to-detect structures highlights an important limitation of the current approach. Reducing annotation requirements for such classes represents an important direction for future work, potentially through semi-supervised or data-efficient learning strategies. This analysis of dense-core granules is presented at the end of the Results section.

### Overview of thrombosis dataset

To highlight the biological applicability of our 3D analysis pipelines, we present one of the most comprehensive 3D ultrastructural analyses of platelet activation to date, encompassing thousands of platelets and their organelles. Our primary objective was to infer organelle secretion dynamics during thrombosis, a key feature of platelet activation with critical roles in hemostasis and thrombo-inflammatory disease progression.

We take examples from four conditions representing increasing degrees of platelet activation. Resting platelets, isolated directly from circulation and immediately fixed, served as the physiological baseline **(Figure 6A)**. The remaining three conditions consist of platelets imaged from thrombi induced *in vivo* using the ferric chloride (FeCl₃) injury model in mouse carotid arteries. This chemical model produces endothelial damage via oxidative stress, triggering localized platelet adhesion, an early step in platelet activation leading to thrombus formation [Grover and Mackaman, 2020]. Varying FeCl₃ concentrations (5.5%, 6%, and 10%) produced thrombi of differing severity as indicated by electron dense iron accumulation, capturing a spectrum of platelet activation states, microenvironmental signaling conditions, and structural variability associated with increasing endothelial injury. The selected ferric chloride concentrations (5.5%, 6%, and 10%) were chosen to span a physiologically and experimentally relevant range. Lower concentrations (5.5% and 6%) produce less severe endothelial injury and are thought to better approximate thrombotic conditions relevant to human disease, where occlusive clotting is less extreme; the inclusion of two closely spaced low concentrations allows finer resolution of early activation states. In contrast, 10% FeCl₃ represents a higher-intensity condition that is widely used in the literature to reliably induce robust thrombus formation [Eckly et al., 2011]. Together, these concentrations enable sampling across a spectrum of injury severity and platelet activation states. At each concentration. samples from two mice were analyzed qualitatively as non-isotropic image stacks (150 nm XY, 1 μm in Z) by SBF-SEM and showed comparable 2D outcomes. For each sample, individual planes spaced 100 microns apart were montage imaged at 10 nm XY pixel size. Again, results for the paired samples at each concentration were similar. The choice of samples for detailed isotropic SBF-SEM imaging was random.

**Fig. 6.**
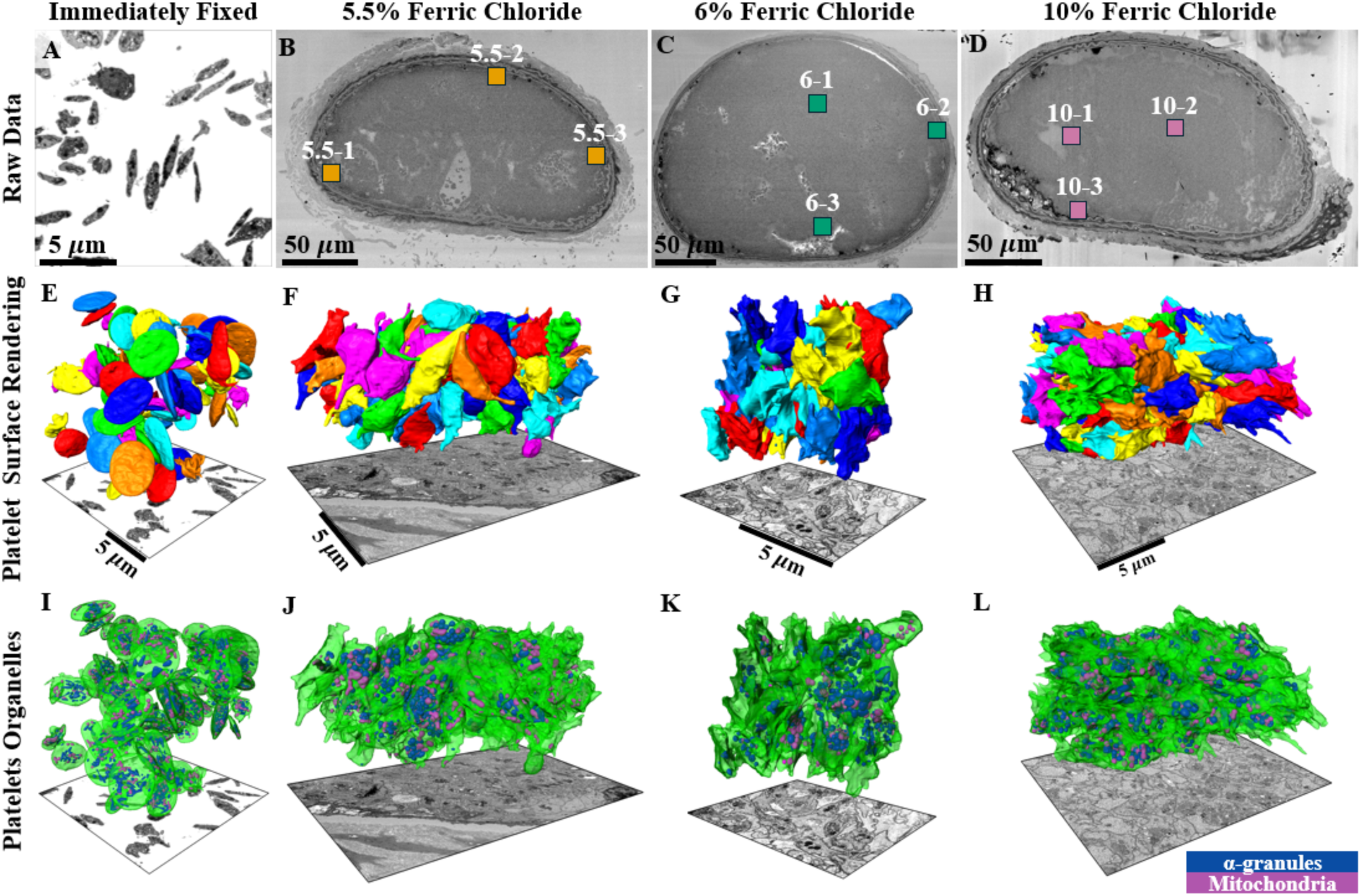
Overview of the dataset. **(A)** Electron micrograph of immediately fixed platelets from a blood suspension, serving as the control. **(B to D)** Ortho-slices through the mid-depth of three thrombi within mouse carotid arteries, formed by applying varying concentrations of ferric chloride. Locations of regions of interests (ROIs) within the thrombi are indicated by rectangle markers. **(E to H)** 3D surface renderings of segmented cells from select ROIs, where each color represents a unique cell (colors may be reused). Cells intersecting the dataset borders were excluded to ensure that only fully captured cells were retained for quantification. **(E)** Segmented cells from the entire immediately fixed dataset. **(F)** Segmented cells from ROI 5.5-2. **(G)** Segmented cells from ROI 6-3. **(H)** Segmented cells from ROI 10-1. **(I to L)** Segmented platelets from the previous row shown with translucent green membranes and internal organelles, with α-granules in blue and mitochondria in magenta. Note that of the platelets present in J-L, not all appeared to contain abundant α-granules and mitochondria. This was true whether the ROI was peripherally located in the thrombus or more centrally located.

Mid-thrombus regions (ROIs), 40 μm x 40 μm x 40 μm were selected for in-depth, high resolution FIB-SEM imaging **Figure 6B–D)**. The 3D TIFF stacks with near-isotropic voxel size (5 nm × 5 nm × 7 nm) were used for the detailed analysis of individual platelets and their subcellular components. ROIs were deliberately spread throughout the thrombus to capture spatial heterogeneity within the clot microenvironment, as indicated by the colored boxes in **Figure 6B–D**. No thrombus cross-section is shown for the control platelets **(Figure 6A)**, as this dataset captures all resting, unactivated, non-adherent platelets, unlike the thrombus samples where ROIs were selected for detailed imaging.

To extract quantitative information, we applied our tailored deep learning-based segmentation pipeline. Surface renderings of segmented platelets from select ROIs are shown in the second row of **Figure 6 (E–H)**, with each color representing a unique platelet (some colors may be reused). To visualize subcellular architecture, segmented platelets were rendered with translucent membranes and organelles highlighted (α-granules in blue, mitochondria in magenta; **Figure 6I–L**).

This dataset enables direct comparison of the ultrastructural characteristics of resting versus thrombus-associated platelets, including differences in organelle content such as α-granules, dense granules, and mitochondria. These organelles play distinct roles in hemostasis, coagulation, and inflammatory signaling, and the spatial resolution and statistical power of our dataset provide a unique opportunity to dissect single-cell morphological changes across varied thrombotic environments. Qualitative observations indicate substantial platelet-to-platelet variability in α-granule and mitochondrial content within induced thrombi, in contrast to the relative uniformity observed in resting platelets.

### Quantitative analyses of organelle distribution

Detailed quantitative analyses are critical for understanding organelle distribution and degranulation behavior at single-cell resolution. In **Figure 7**, we present frequency histograms quantifying the number of organelles per platelet across all experimental conditions. Each point represents a single platelet, with its y-axis position reflecting the absolute number of mitochondria, condensed α-granules, or decondensed α-granules within that cell. Decondensed α-granules arise from activated α-granules that fail to fully collapse into the plasma membrane during the process of content release [12, 13].

**Fig. 7.**
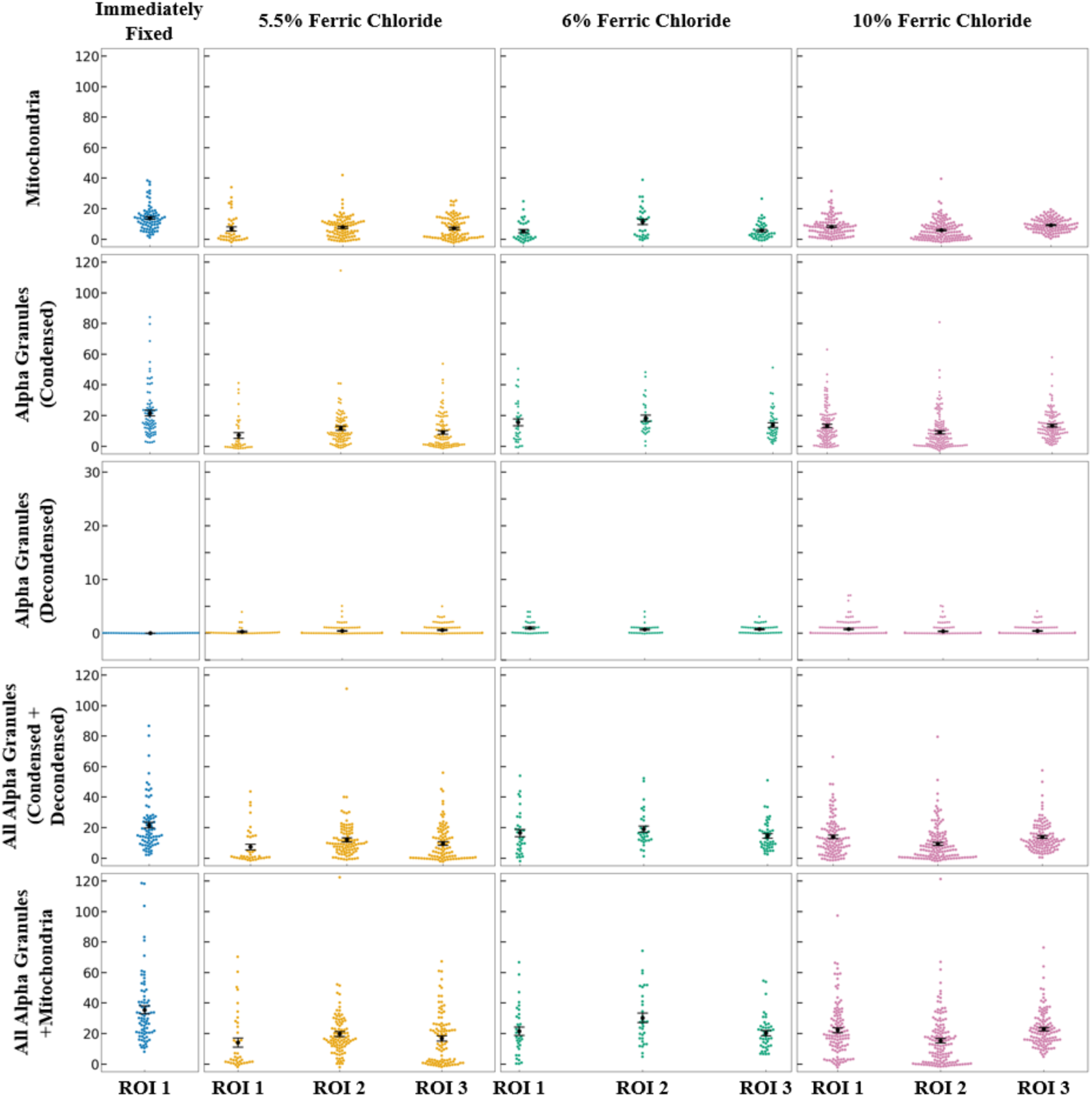
Organelle counting. Each point represents a single platelet, with the y-axis showing the number of mitochondria (top row), condensed α-granules (second row), decondensed α-granules (third row), both condensed and decondensed α-granules (fourth row), or all combined (bottom row) in each cell. Data are shown across induction conditions: Immediately Fixed (leftmost column) represents the control group, while 5.5%, 6%, and 10% indicate increasing concentrations of ferric chloride used to induce clot formation. Points at zero on the bottom row indicate fully degranulated platelets. Error bars represent the standard error of the mean across cells.

We find that mitochondria and α-granules remain consistently abundant: most platelets contain approximately 15 mitochondria and 20 α-granules, regardless of ferric chloride concentration. Interestingly, our data also reveals the presence of fully degranulated platelets, or cells devoid of all major organelle types. These cells are present across all ROIs but are particularly prevalent in the peripherally located ROI 1 and ROI 3 from the 5.5% ferric chloride group, and the more centrally located ROI 2 from the 10% group.

This phenomenon of complete platelet degranulation can further be seen in 2D slices extracted from the FIB-SEM volumes (**Figure 8 A**). In addition, these images illustrate that this process tends to occur in spatially localized regions in which platelets appear uniformly degranulated (**Figure 8 B**), supporting the notion that activation occurs in clustered domains rather than in uniform spatial gradients across the thrombus. Similar patterns have been observed across larger fields of view via SBF-SEM, reinforcing the robustness of this phenomenon. In puncture-wound time-series experiments, similar two-dimensional clusters of degranulated platelets have been observed lining the surfaces of vaulted cavities within the developing thrombus [44,45].

**Fig. 8.**
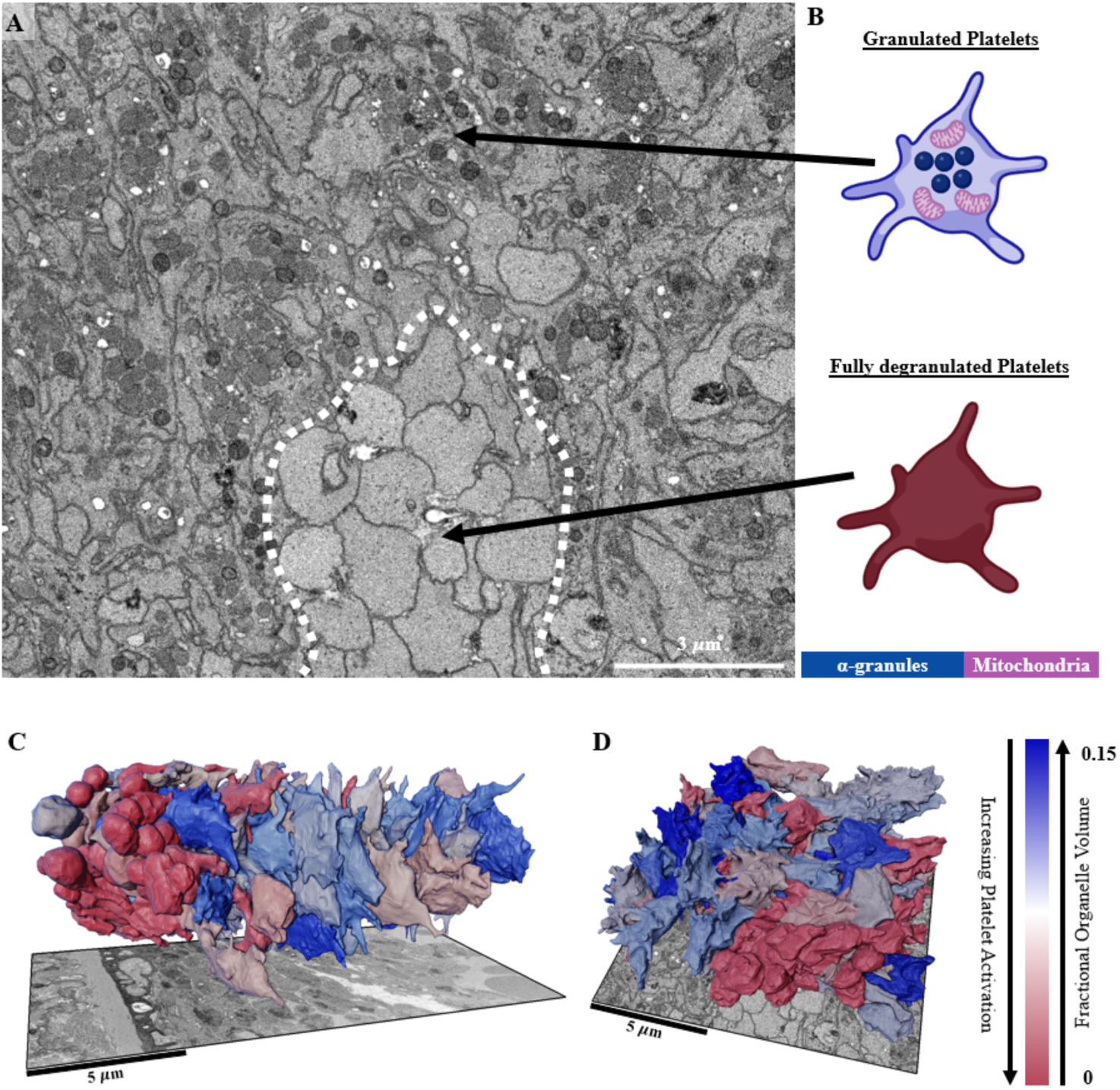
Spatial heterogeneity of platelet degranulation in a thrombus. **(A)** Raw electron micrograph showing a clear pocket of complete platelet degranulation. **(B)** Cartoon showing the organelle contents of platelets in the two adjacent regions, one high granulation (blue) and one completely degranulated (red). α-granules are shown as blue spheres and mitochondria as magenta bean-shaped structures. **(C-D)** 3D surface renderings of segmented platelets with heat map showing the degree of degranulation for each platelet, with red indicating highly degranulated (activated) platelets and blue indicating granulated (less activated) platelets. Degranulation occurs in spatially clustered regions, consistent with the short-range nature of platelet activation signaling mechanisms.

To further characterize the spatial distribution of platelet activation, we generated single-cell resolution heat maps (**Figure 8 C-D**), depicting the degree of granule abundance in each segmented platelet. In these maps, red indicates highly degranulated platelets, whereas blue represents granule-rich, less activated cells. This analysis reveals discrete microdomains within the thrombus, where clusters of degranulated platelets are surrounded by less activated neighbors.

Such observed heterogeneity in activation state is consistent with models of localized, short-range signaling within thrombi, where tightly packed platelets may undergo differential activation in response to local biochemical cues. Importantly, such spatial patterns would be entirely obscured using traditional bulk biochemical assays, which average measurements across large, heterogeneous cell populations. These results underscore the critical value of high-resolution, spatially resolved volumetric imaging for dissecting the complex and heterogeneous nature of platelet activation within the thrombus microenvironment and validate results from previous 2D analysis [Rhee et al., 2021; Rhee et al., 2026].

Strikingly, our analysis also reveals a bimodal, or binary, pattern of organelle retention. Most platelets appear to fall into one of two categories: either nearly fully granule-rich or completely degranulated. Intermediate states are rare. Such a pattern implies a threshold-dependent signaling mechanism governing organelle release, a hypothesis that is suggested by prior 2D studies indicating localized degranulation zones [Pokrovskaya et al., 2025]. The absence of α-granules and mitochondria appears to be correlated, with many platelets lacking one also lacking the other. The mechanistic basis for this pattern is currently unknown and will require further investigation.

To further probe the nature of this binary depletion, we excluded fully degranulated platelets from our analysis and examined only those cells with nonzero organelle content. When comparing the mean fractional organelle volumes of these granule-containing platelets across all samples, as shown in **Figure 9**, we observed no statistically significant differences. These findings provide quantitative evidence that most platelets do not secrete α-granules, and that mitochondrial content remains largely unchanged, consistent with their role as bioenergetic organelles. Importantly, when α-granule release is assessed based on the presence of decondensed α-granules, only a small fraction of platelets exhibited detectable secretion, even at high ferric chloride concentrations. Notably, degranulation appears to occur in an all-or-none fashion, largely independent of the overall level of endothelial injury. These data support a model in which α-granules and mitochondria respond to local rather than global signaling, distinct from the more pronounced release behavior of dense granules, as explored in the following section.

**Fig. 9.**
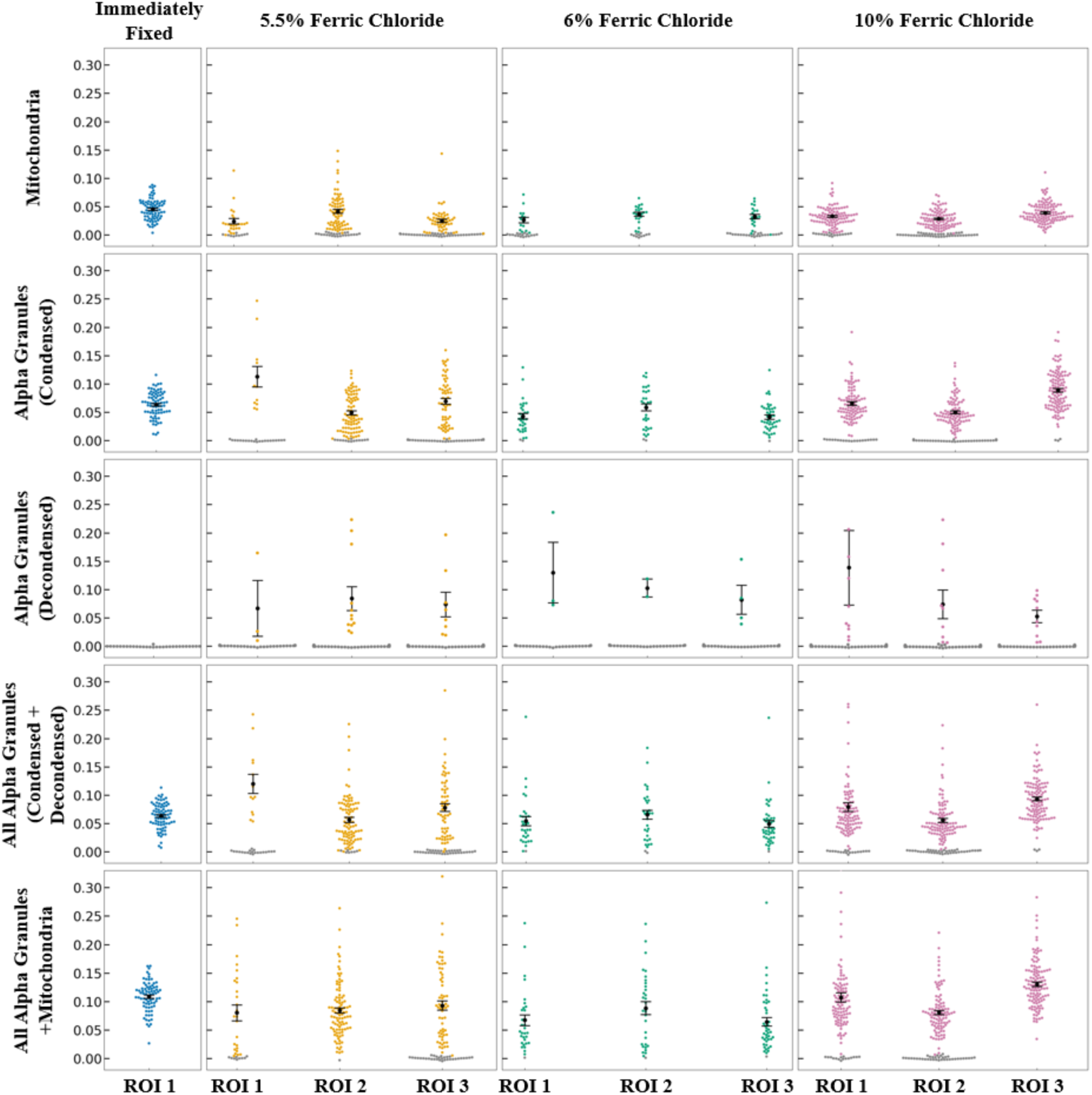
Organelle fraction. Each point represents a single platelet, with the y-axis showing the fraction of a platelet’s volume occupied by mitochondria (top row), condensed α-granules (second row), decondensed α-granules (third row), both condensed and decondensed α-granules (fourth row), or all combined (bottom row). The volume fraction was used instead of organelle counts to minimize systematic error and provide a more stable measure of organelle content. Data are shown across fixation conditions: Immediately Fixed (leftmost column) represents the control group, while 5.5%, 6%, and 10% indicate increasing concentrations of ferric chloride used to induce clot formation. Data from all three ROIs within each injury condition were combined into a single group for statistical comparison. Among platelets retaining organelles, no significant differences in fractional organelle volume (combined volume of all three organelle types, pooled across ROIs) were observed across injury conditions (Kruskal–Wallis H = 2.91, p = 0.088), indicating little to no collapse of organelle membranes into the platelet plasma membrane during the formation of highly adherent, platelet-rich occlusive thrombi. Grey points indicate cells lacking the organelle of interest and were excluded from statistical comparisons and error bar calculations.

A quantitative summary of platelet α-granule content across conditions is provided in **Table IV**. Consistent with the analysis in **Figure 9**, only a minority of platelets exhibited evidence of α-granule release, as indicated by the presence of decondensed α-granules, whereas the majority retained their α-granule content across all conditions. These data provide a population-level view of the heterogeneity in platelet activation and support the conclusion that α-granule secretion occurs in a limited subset of platelets within occlusive thrombi.

**Table IV:**
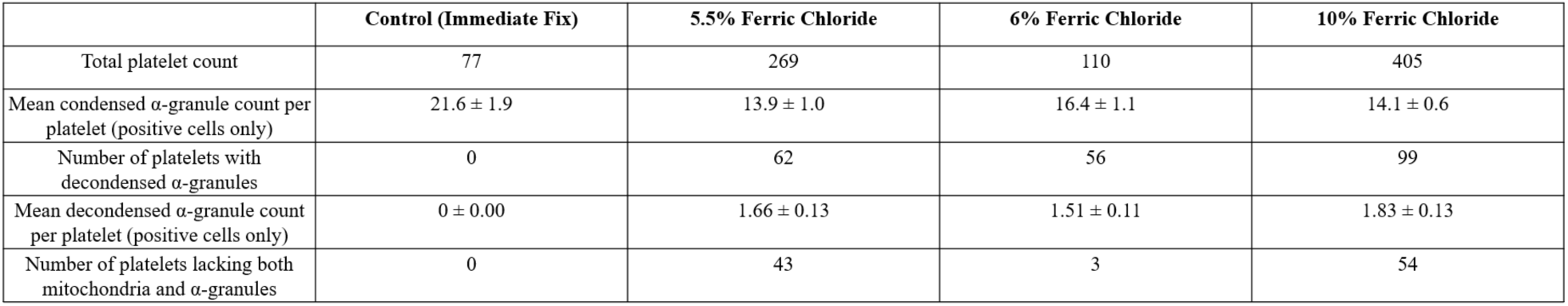
Quantitative summary of platelet and organelle metrics across thrombotic conditions.

### Dense granule behavior

Building on our observations of α-granules and mitochondria, we next examined dense granules. Due to the low abundance of dense granules and challenges in predicting sparse classes using our machine learning models, we opted to perform simple manual counting of dense granules (**Fig.** 10, A–C).

**Fig. 10.**
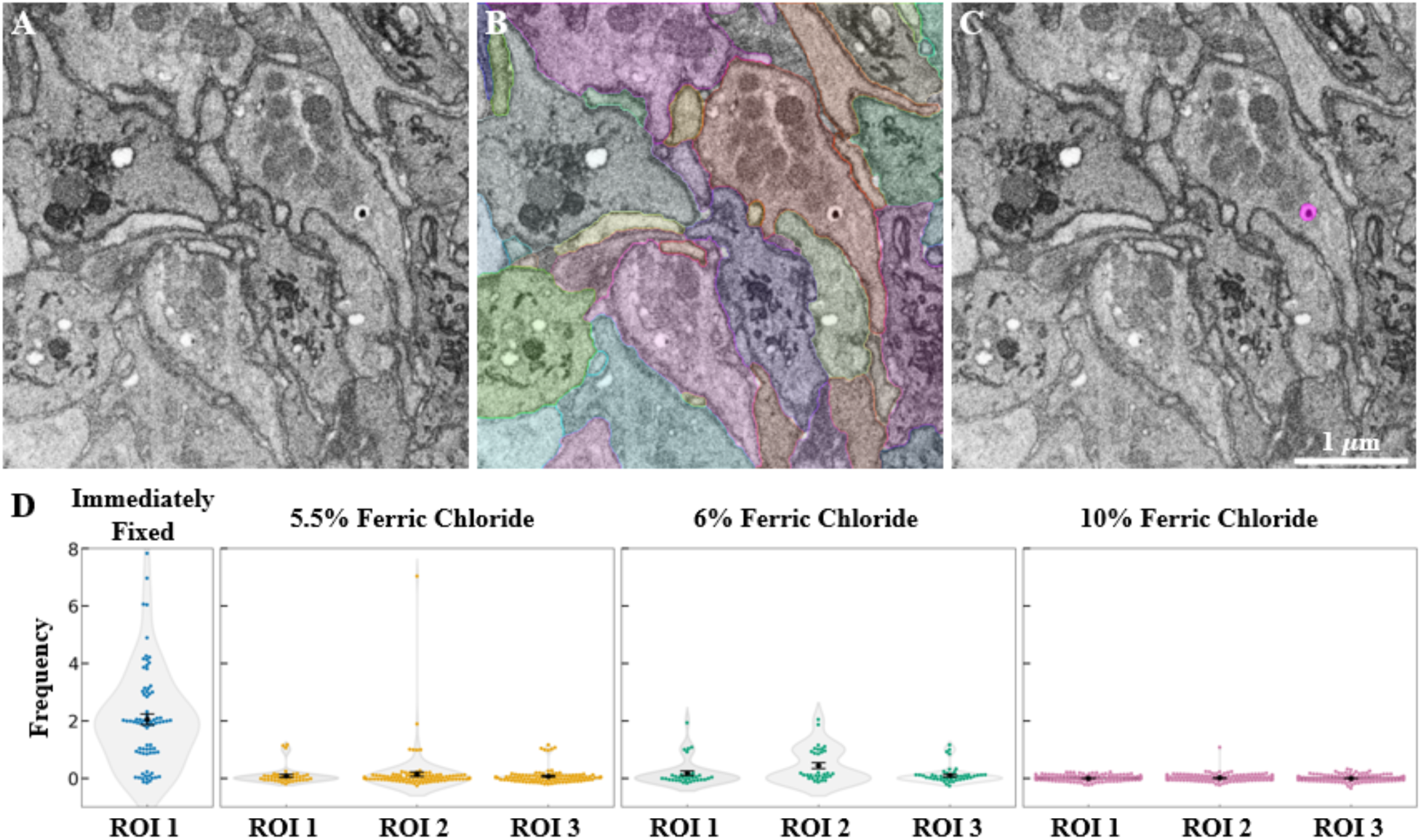
Dense granule identification and release patterns. Electron micrographs showing **(A)** a raw EM image, **(B)** segmented platelets (each color represents a unique platelet), and **(C)** a representative example of a dense core granule (in pink). **(D)** Each point represents a single platelet, with the y-axis indicating the number of dense core granules occupying each cell. Data are shown across fixation conditions: Immediately Fixed (leftmost column) represents the control group, while 5.5%, 6%, and 10% correspond to increasing concentrations of ferric chloride used to induce clot formation. Error bars represent the standard error of the mean across cells.

We observed that most platelets within thrombi exhibit near-complete loss of dense granules. This consistent depletion aligns with the well-established role of dense granules as the first organelle type secreted during platelet activation and integrin-dependent platelet adhesion [Yadav and Storrie, 2017], preceding the later release of α-granule contents and mitochondria [Pokrovskaya et al., 2020].

Comparing across organelle types, we observed a striking difference in depletion patterns. While overall organelle counts were reduced in thrombus-associated platelets compared to resting platelets, reflecting secretion upon activation, this depletion was highly organelle-specific. Dense granules exhibited uniform and substantial loss across all thrombotic conditions, underscoring their function as high-intensity, early responders in the platelet activation cascade **(Fig. 10, D)**. In contrast, α-granules and mitochondria showed no significant change in their average numbers, despite their known involvement in later-stage coagulation [Kim et al., 2020] and inflammatory signaling.

Overall, these results suggest that organelle release is governed by distinct regulatory mechanisms. Dense granules undergo a rapid, near-global response, while α-granules and mitochondria follow a more binary, spatially patterned depletion, likely driven by localized signaling domains within the thrombus. While this analysis focuses on secretion-related changes in organelle content, platelet function in hemostasis is also critically dependent on adhesion and aggregation. The observed uniform depletion of dense granules may reflect their role in rapidly amplifying platelet adhesion responses, whereas the more stable α-granule counts suggest that their contribution may not be fully captured by abundance alone. Integrating structural and adhesion-related metrics will be an important direction for future work.

### Limitations of the Study

Despite providing very high-resolution volumetric insights into platelet ultrastructure, this study is constrained by several factors. First, the FIB-SEM volumes analyzed represent a small fraction of the overall clot (typically 40 μm × 40 μm × 40 µm per ROI). Although multiple regions were sampled across each thrombus, all were collected at similar depths near the clot midpoint, the first formed part of the clot, limiting our ability to assess spatial organization through an entire thrombus. This limitation arises primarily from the time required for both FIB-SEM data acquisition and analysis. We are currently addressing it through correlative approaches using high-resolution SBF-SEM mosaics to generate broader, multiscale maps of clot architecture [Faruque et al. 2025]. Second, the study focuses exclusively on thrombi induced by ferric chloride occlusion, which may differ from other vascular injury models; ongoing work with a puncture-wound model aims to test the generality of these findings. Finally, the imaging captures static snapshots of fixed samples, precluding direct observation of temporal events such as degranulation dynamics or mitochondrial remodeling.

## Discussion

In this study, we present a high-throughput framework for the segmentation and analysis of platelet morphology and organelle fates during thrombosis using volumetric electron microscopy (vEM) combined with deep learning. By leveraging isotropic FIB-SEM imaging and a multi-axis, nnU-Net–based segmentation pipeline, we achieved robust, instance-level detection of platelets, α-granules, mitochondria, and dense granules across thrombi formed under varying occlusion induction intensities demonstrating the extended biological analysis capabilities made feasible by our pipelines.

Methodologically, our segmentation framework demonstrates that high-quality volumetric results can be obtained using 2D neural networks by aggregating predictions from three orthogonal axes. This strategy captures 3D context without the computational cost or extensive training data required for fully 3D architectures. By eliminating the need for specialized computing resources, our pipeline enables large-scale, quantitative ultrastructural analysis on standard desktop hardware, making volumetric studies broadly accessible. Importantly, our approach succeeds in the crowded environment of platelet-rich thrombi, where cell boundaries are tightly apposed and membranes are frequently shared between neighboring cells. Such densely packed ultrastructure presents far greater challenges than segmenting isolated cells, where many existing auto-segmentation approaches suffice. While applied here to platelets within thrombi, this framework is readily adaptable to other cellular systems and pathophysiological processes involving subcellular remodeling, including cancer, inflammation, and neurodegeneration.

Our large-scale, single-cell methods application to platelet biology reveals previously unobservable features of thrombus biology. Most notably, we identified striking, organelle-specific depletion patterns in activated platelets. Some platelets exhibited partial release of α-granule contents, as indicated by the presence of decondensed α-granules, a residual structure following content release. Within distinct three-dimensional clusters of platelets, α-granules and mitochondria displayed coordinated binary (all-or-none) loss. In contrast, dense granules showed a uniform and near-global depletion across all thrombotic conditions. These patterns of α-granule and mitochondrial co-loss were spatially clustered at mere micron scales, reflecting highly localized signaling events within thrombi. Such spatial organization can only be inferred by traditional 2D assays and not at all in population-averaged assays, underscoring the power of volumetric, spatially resolved imaging in uncovering emergent behaviors in multicellular systems. Dot-plot analysis quantitatively revealed substantial heterogeneity, with many platelets completely lacking major organelles, highlighting the diverse activation states within individual thrombi.

Previous investigations of platelet activation and organelle dynamics have largely relied on two-dimensional imaging and automated scoring methods. While informative, these approaches are limited in capturing volumetric context, spatial heterogeneity, and full organelle distributions. For example, 2D analyses suggested graded “shaded” α-granule depletion and highly localized mitochondria loss, but the planar perspective precluded definitive quantification of 3D organelle depopulation. In contrast, our isotropic 3D imaging and segmentation pipeline provides a complete volumetric view, enabling rigorous assessment of platelet activation states, spatial clustering, and heterogeneity. This approach not only validates and extends prior 2D observations but also reveals new insights, such as the short-range, local nature of activation and the striking organelle-specific responses and validates inferences from 2D studies [Rhee et al. 2026; Pokrovskaya et al., 2025]. Consistent with prior 2D kinetic studies of puncture-induced thrombus formation, which report dynamic exposure of degranulated platelet clusters at the thrombus surface, our findings provide a three-dimensional framework for interpreting these phenomena. Extending this approach to time-resolved 3D analysis of thrombus formation is expected to yield further mechanistic insight.

Overall, our findings provide strong evidence for organelle-specific regulatory mechanisms during platelet activation. Dense granules act as rapid, high-intensity early responders, exhibiting uniform depletion across thrombotic conditions, while α-granules and mitochondria follow a more binary, spatially patterned release, likely governed by localized signaling domains. The discovery that platelet activation and organelle depletion occur over very short distances, just a few microns, highlights the importance of considering spatial heterogeneity in thrombus biology. By enabling quantitative, high-resolution volumetric analysis, our work sets a new standard for studying platelet ultrastructure *in situ* and offers a scalable platform for future investigations into cellular microenvironments and emergent multicellular behaviors.

## Data and materials availability

Code can be found as a lightweight Jupyter notebook for initializing and running the neural network at our public GitHub repository: https://github.com/samuel-fulton/Cell-Seg. Data examples can be found also at our public OSF repository: https://osf.io/mpysc/overview.

## Acknowledgments

We thank Joshua Kim and Denzel Cruz for collecting portions of the FIB-SEM data; Zijie (Jed) Yang for early conceptual contributions that inspired the multi-axis pipeline design; Elizabeth R. Driehaus and Sidney W. Whiteheart for performing the surgical induction and excision of thrombi; and members of the Storrie laboratory for insightful biological discussions that motivated this project. This work utilized the computational resources of the NIH HPC Biowulf cluster (http://hpc.nih.gov). Platelet cartoons created in BioRender.

## Author contributions

Conceptualization: SJF, CMB, BS

Methodology: SJF, CMB

Investigation: SJF, CMB

Visualization: SJF, CMB

Supervision: MAA, BS, RDL

Writing—original draft: SJF, CMB

Writing—review & editing: SJF, CMB, MAA, BS, RDL

## AI Use Declaration

The authors used OpenAI’s ChatGPT for general code debugging during the development of the U-Net neural network. Standard tools for grammar checking (e.g., Grammarly) were employed to the manuscript, without contributing to the scientific content. All scientific content, data analysis, and interpretations were created and verified by the authors.

